# A rose flavor compound activating the NRF2 pathway in dendritic cells ameliorates contact hypersensitivity in mice

**DOI:** 10.1101/2022.02.08.479651

**Authors:** Naoki Kodama, Hikaru Okada, Masakazu Hachisu, Miki Ando, Naoto Ito, Kazuki Nagata, Yayoi Yasuda, Ikumi Hiroki, Takuya Yashiro, Gaku Ichihara, Masayuki Yamamoto, Chiharu Nishiyama

## Abstract

Dendritic cells (DCs), which are typical antigen-presenting cells, localize to various sites in the body, particularly the front line of infection as sentinels, and are involved in innate and adaptive immune responses. Although the functions of DCs, such as pathogen-induced cytokine production and antigen-specific T cell activation, are important for host defenses against infection and tumorigenesis, the hyper- and/or extended activation of DCs leads to inflammatory and autoimmune diseases. In the present study, β-damascone, a major ingredient of rose fragrance, was selected from an aroma library as a candidate compound that suppresses antigen-induced immune responses. β-Damascone inhibited the functions of DCs, including the antigen-dependent proliferation of T cells, DC-induced Th1 development, and the TLR ligand-induced production of inflammatory cytokines by DCs. The β-damascone treatment also increased the protein level of the transcription factor NRF2, which plays key roles in antioxidant responses, and the transcription of *Hmox1*, a target gene of NRF2, in DCs. *Nrf2*^−/−^ DCs induced Th1-development and produced large amount of IL-12p40 even in the presence of β-damascone, whereas these functions by *Nrf2*^+/−^ DCs were inhibited by β-damascone under the same conditions. The intake of β-damascone suppressed ear swelling in contact hypersensitivity (CHS) model mice, but not in CHS-induced *Nrf2*^−/−^ mice. Collectively, the present results indicate the potential of the rose aroma compound β-damascone, which suppresses DC-mediated immune responses by activating the NRF2 pathway in DCs, for the prevention and/or attenuation of immune-mediated diseases.

## Introduction

Dendritic cells (DCs), the most typical antigen presenting cells (APCs), contribute to both innate and adaptive immunity. The activation of DCs by pathogens leads to the release of cytokines and chemokines from DCs, and the DC-mediated expansion of antigen (Ag)-specific T cell clones. Therefore, DC activation is essential for host defenses against infection; however, the hyper- and/or extended activation of DC-mediated immune responses causes inflammatory and autoimmune diseases.

Humans have traditionally obtained pharmacological compounds from natural products derived from plants and bacteria. A number of studies have examined the immunomodulatory effects of natural compounds, particularly phytochemicals, which are produced in plants to resist various stresses, including UV and insect pests (Mainardi, Kapoor and Bielory 2009). Resveratrol is the most well-known polyphenolic stilbenoid (Baur et al. 2006, Jang et al. 1997) and has been shown to prevent inflammation-related diseases by suppressing the TLR signaling-induced expression of proinflammatory genes (Malaguarnera 2019). Regarding scent phytochemicals, several food ingredients, including curcumin (Kunnumakkara et al. 2017) and menthol (Oz et al. 2017), are expected to possess therapeutic potential.

In the present study, we performed screening to identify novel immunomodulators and selected β-damascone, a major ingredient of rose fragrance, from approximately 150 types of aroma compounds. We then investigated the molecular mechanisms by which β-damascone regulates Ag-dependent T cell proliferation. The obtained results showed that β-damascone suppressed DC-mediated immune responses and revealed that its activation of the transcription factor NF-E2-related factor 2 (NRF2) pathway in DCs was one of the underlying causes. We then confirmed the effects of β-damascone on immune responses *in vivo* using a contact hypersensitivity (CHS) mouse model. Taken together, the present results indicate that β-damascone, which suppresses DC-related immune responses, contributes to the prevention and/or attenuation of immune-mediated diseases.

## Materials and Methods

### Mice and cells

C57BL/6 and Balb/c mice were purchased from Japan SLC (Hamamatsu, Japan), and OT-II mice were obtained from The Jackson Laboratory (USA). *Nrf2*^−/−^ mice were previously generated (Itoh et al. 1997). Mice were maintained under specific pathogen-free conditions. All animal experiments were performed in accordance with the guidelines of the Institutional Review Board of Tokyo University of Science. The current study was specifically approved by the Animal Care and Use Committees of Tokyo University of Science: K21004, K20005, K19006, K18006, K17009, and K17012. Bone marrow-derived DCs (BMDCs) were generated from whole BM cells by cultivation in RPMI-1640-based media supplemented with 20 ng/mL mGM-CSF (BioLegend, San Diego, CA, USA) as previously described (Kanada et al. 2011). Naïve CD4^+^ T cells were isolated from the spleen by using the MojoSort Mouse Naïve CD4^+^ T Cell Isolation Kit (#480040, BioLegend). CD4^+^ T cells isolated from the OT-II spleen were cocultured with BMDCs, which were generated from C57BL/6 mice and were preincubated with OVA peptide 323-339 (POV-3636-PI, Peptide Institute, Inc., Osaka, Japan). For Th1 polarization, 10 ng/mL mIL-12 (PeproTech Inc., Rocky Hill, NJ, USA) and 10 μg/mL anti-IL-4 Ab (clone 11B11, BioLegend) were added to the culture media. Th2 polarization was induced with 20 ng/mL IL-4 (PeproTech) and 10 μg/mL anti-IL-12 Ab (clone C17.8, BioLegend).

β-Damasone (#34059, Vigon International, East Stroudsburg, PA, USA) was diluted with DMSO. LPS (#L3024, Wako), poly-I:C (#P0913, Sigma), R848 (AG-CR1-3582-M005, AdipoGen, Liestal, Switzerland), and CpG (ODN1826, InvivoGen, San Diego, CA, USA) were used to stimulate BMDCs. DAPI (#11034-56, Nacalai Tesque Inc., Kyoto, Japan) was used to determine cell viability.

### Enzyme-linked immunosorbent assay (ELISA)

The concentrations of mouse IL-2, IL-6, TNF-α, and IL-12p40 were determined by ELISA kits purchased from BioLegend (#431004, #431315, #430915, and #431604, respectively).

### Flow cytometry

To analyze the proliferation of T cells, CFSE (eBioscience Inc., San Diego, CA, USA) was used. Cell surface MHC class II and CD86 on BMDCs were stained with anti-I-A/I-E-PerCP (clone M5/114.15.2, BioLegend) and anti-CD86-PE (clone GL-1, BioLegend), respectively. Intracellular IFN-γ and IL-4 were stained with anti-IFN-γ-PE/Cyanine7 (clone XMG1.2, BioLegend), and anti-IL-4-PE (clone 11B11, BioLegend), respectively, with anti-CD4-FITC (clone GK1.5, BioLegend) after treatment with the Fixation Buffer (#420801, BioLegend) and the Intracellular Staining Perm Wash Buffer (#421002, BioLegend). Fluorescence was detected by a MACS Quant Analyzer (Miltenyi Biotech) and analyzed with FlowJo (Tomy Digital Biology, Tokyo, Japan).

### Western blot analysis

Western blot analysis was performed as previously described (Kitamura et al. 2012) with anti-phospho-IkBα (Ser32) (clone 14D4, Cell Signaling), anti-IkBα (clone L35A5, Cell Signaling), anti-NRF2 (clone D1Z9C, Cell Signaling), and anti-β-actin (clone AC-15, Sigma-Aldrich) Abs.

### Quantitative RT-PCR

Total RNA was extracted from BMDCs using the ReliaPrep RNA Cell Miniprep System (#Z6012, Promega, Madison, USA) and from the skin using ISOGEN (#311-02501, Nippongene, Tokyo, Japan). Synthesis of cDNA, and quantitative PCR were performed as previously described (Ito et al. 2021). The nucleotide sequences of the primer sets are listed in **Supplementary Table SI**.

### CHS model

Mice were sensitized on shaved abdominal skin with 25 μl 0.5% (w/v) DNFB in acetone/olive oil (4:1) and were challenged with an application of 20 μl 0.25% DNFB on the ear at 5 days after sensitization. Ear thickness was measured by a caliper.

### Statistical analysis

To compare two samples, a two-tailed Student’s t-test was used. To compare more than three samples, one-way ANOVA-followed by Tukey’s multiple comparison test or Dunnett’s multiple comparison test was used. *P* values < 0.05 were considered significant.

## Results

### β-damascone was identified as an immunomodulator that suppresses DC-mediated T cell activation

To identify novel immunomodulators, we performed 2-step screening using an aroma-compound library (**Supplementary Table SII**). In the 1st screening, in which APC-dependent T cell proliferation was assayed, 20 candidates were selected from approximately 150 compounds as immunosuppressors. In the 2nd screening, we investigated the inhibitory effects of the candidate compounds that passed through the 1st screening on IL-2 release from OVA-pulsed OT-II splenocytes, and selected β-damascone (**Figure 1A**), a major component of rose aromas, as the most effective inhibitor (**Supplementary Figure S1**). β-Damascone suppressed OVA-induced IL-2 production by OT-II splenocytes (**Figure 1B**) and the APC-dependent proliferation of CD4^+^ T cells (**Figure 1C**) in a dose-dependent manner. Furthermore, β-damascone significantly suppressed the development of Th1 cells induced by OVA-pulsed DCs (**Figure 1D**), but not Th2 cells (**Figure 1E**).

These results indicate that β-damascone suppressed the APC-dependent activation of CD4^+^ T cells and the development of Th1 cells.

**Figure 1.**
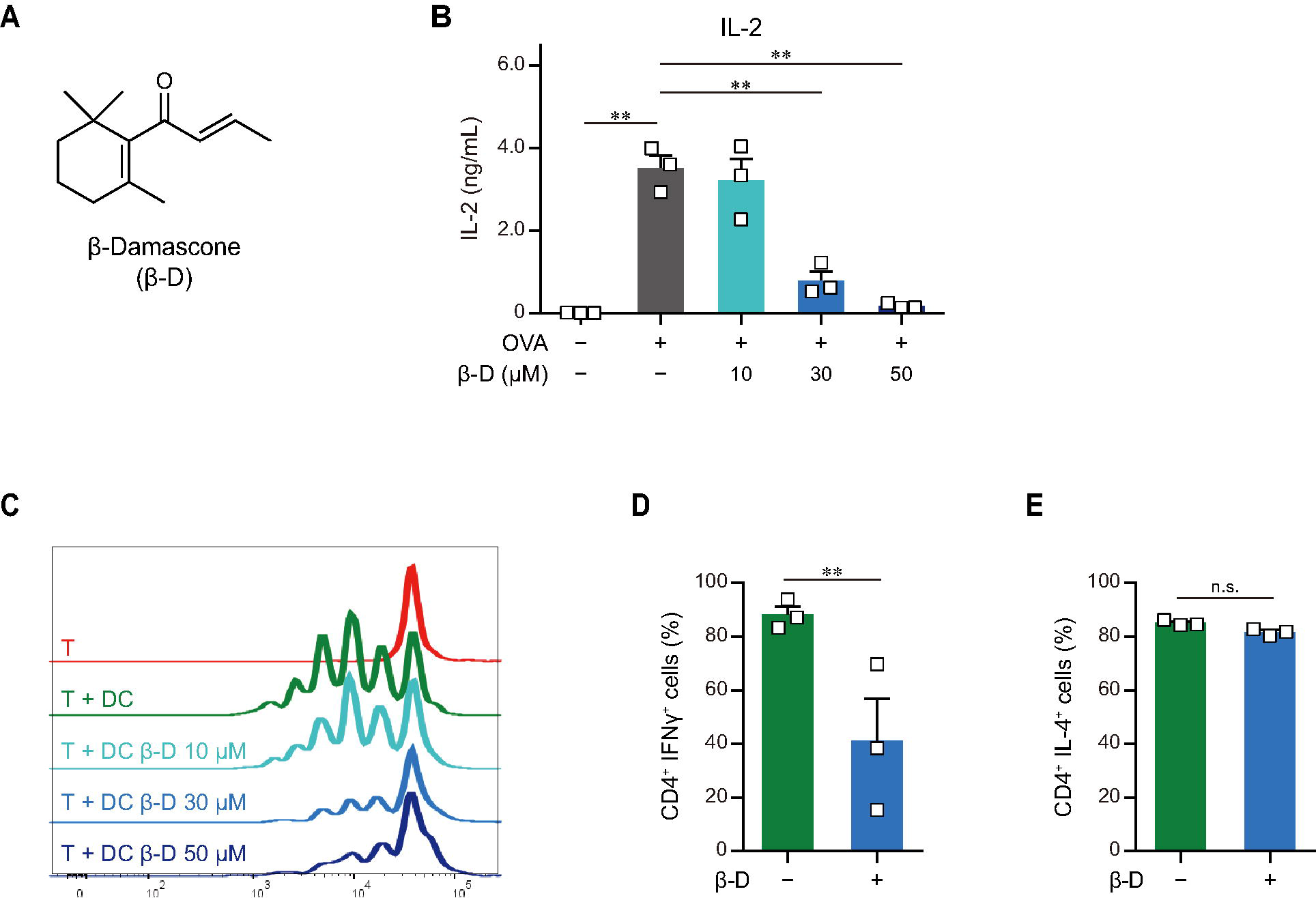
Effects of β-damascone on the DC-mediated activation of T cells. **A**. Structure of β-damascone. **B**. IL-2 concentrations in the culture media of OT-II splenocytes. A total of 1.0 × 10^5^ cells obtained from the whole spleen of OT-II mice were maintained in the presence or absence of 500 ng of the OVA peptide and the indicated concentration of β-damascone (β-D) in 200 μL pf culture media for 72 h. **C**. Division of OT-II CD4^+^ T cells cocultured with OVA-pulsed BMDCs. CFSE-labeled OT-II CD4^+^ T cells (5.0 × 10^4^) were cocultured with or without C57BL/6 BMDCs (1.0 × 10^4^), which were pretreated with 500 ng of the OVA peptide, in the presence or absence of the indicated concentrations of β-damascone in 200 μL of culture media for 72 h. **D**. Effects of β-damascone on the frequencies of Th1 and Th2 cells that developed under polarizing conditions in a DC-dependent manner. OT-II naïve CD4^+^ T cells were cocultured with OVA-pulsed C57BL/6 BMDCs under Th1- or Th2-polarizing conditions in the presence or absence of β-damascone (30 μM) for 72 h. The Tukey-Kramer test was used (**B**, **C**, **D** and **E**). *; *p* < 0.05, **; *p* < 0/01, n.s.; not significant.

### β-damascone suppressed the LPS-induced activation of DCs

We investigated the effects of β-damascone on DC functions by analyzing the expression of cytokines and the cell surface expression levels of APC-related molecules in LPS-stimulated BMDCs. As shown in **Figure 2A**, β-damascone appeared to reduce the mRNA levels of IL-6, IL-12p40, and TNF-α in DCs, which were increased by the LPS stimulation, in a dose-dependent manner at concentrations that were markedly lower than those exhibiting cytotoxicity (**Supplementary Figure S2**). The concentrations of these inflammatory cytokines, particularly IL-6 and IL-12p40, in the culture media of LPS-stimulated DCs were also decreased in the presence of β-damascone (**Figure 2B**), suggesting that the suppression of the LPS-induced transactivation of cytokine genes by β-damascone was reflected in protein release. Furthermore, the flow cytometric analysis revealed that the LPS-induced up-regulation of MHC class II and CD86 was inhibited in β-damascone-treated DCs (**Figure 2C**). We confirmed that the increased phosphorylation of IκB in LPS-stimulated DCs was reduced by the pretreatment with β-damascone (**Figure 2D**).

**Figure 2.**
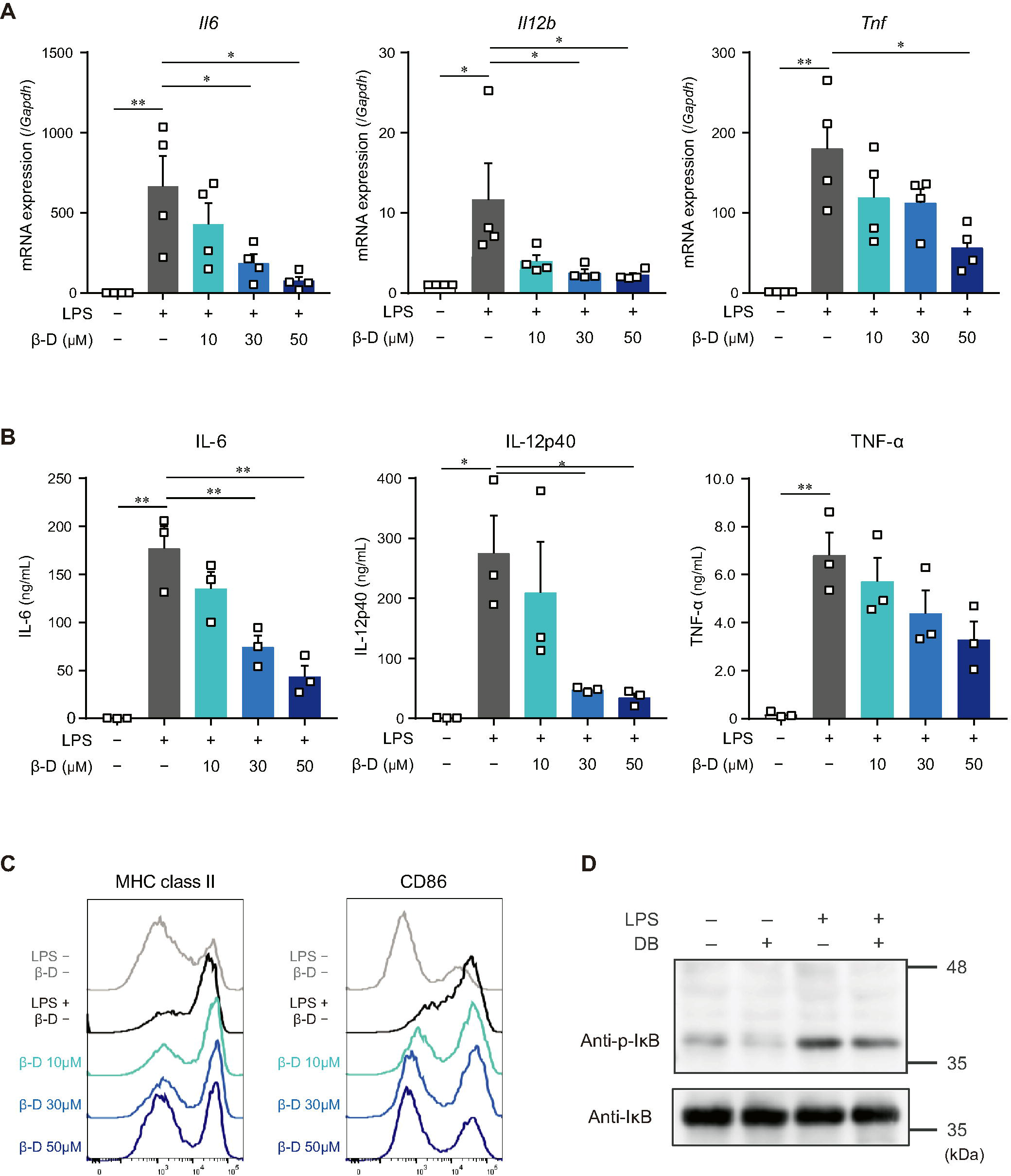
β-Damascone suppressed the LPS-induced activation of DCs. **A**. Messenger RNA expression levels in BMDCs. **B**. Cytokine concentrations in the culture media of BMDCs. **C**. Cell surface expression levels of MHC class II and CD86 on BMDCs. **D**. Western blot profiles of phosphorylated IκB and total IκB. BMDCs pretreated with indicated concentrations of β-damascone for 24 h were stimulated with 100 ng/mL LPS. Cells were harvested 3 and 24 h after the LPS stimulation to measure mRNA levels (**A**) and cell surface protein expression levels (**C**), respectively, and culture media were collected at 24 h after the stimulation to assess cytokine concentrations (**B**). In the Western blot analysis, cells were lysed 1 h after the LPS stimulation, and aliquots containing 10 μg of protein were added to each lane (**D**). Data represent the mean ± SEM of three independent experiments performed in triplicate (**A** and **B**). A gating strategy of the flow cytometric analysis of BMDCs is shown in **Supplementary Figure S3**. The Tukey-Kramer test (**A** and **B**) was used. *; *p* < 0.05, **; *p* < 0/01.

These results indicate that β-damascone exerted suppressive effects on LPS-induced inflammatory responses and Ag-presentation activity in DCs.

### Effects of β-damascone on the activation of DCs via other TLRs

The effects of β-damascone on the activation of DCs caused by stimulants other than LPS were examined by using polyI:C, R-848, and CpG. The quantification of mRNA levels (**Figure 3A-3C**) and measurement of protein concentrations (**Figure 3D-3F**) revealed that the transactivation and subsequent protein production of IL-6, IL-12p40, TNF-α, and IL-23p19 in BMDCs, which were induced by a stimulation via TLR3 (**Figure 3A, D**), TLR7/8 (**Figure 3B, E**), or TLR9 (**Figure 3C, F**), were significantly inhibited by β-damascone.

**Figure 3.**
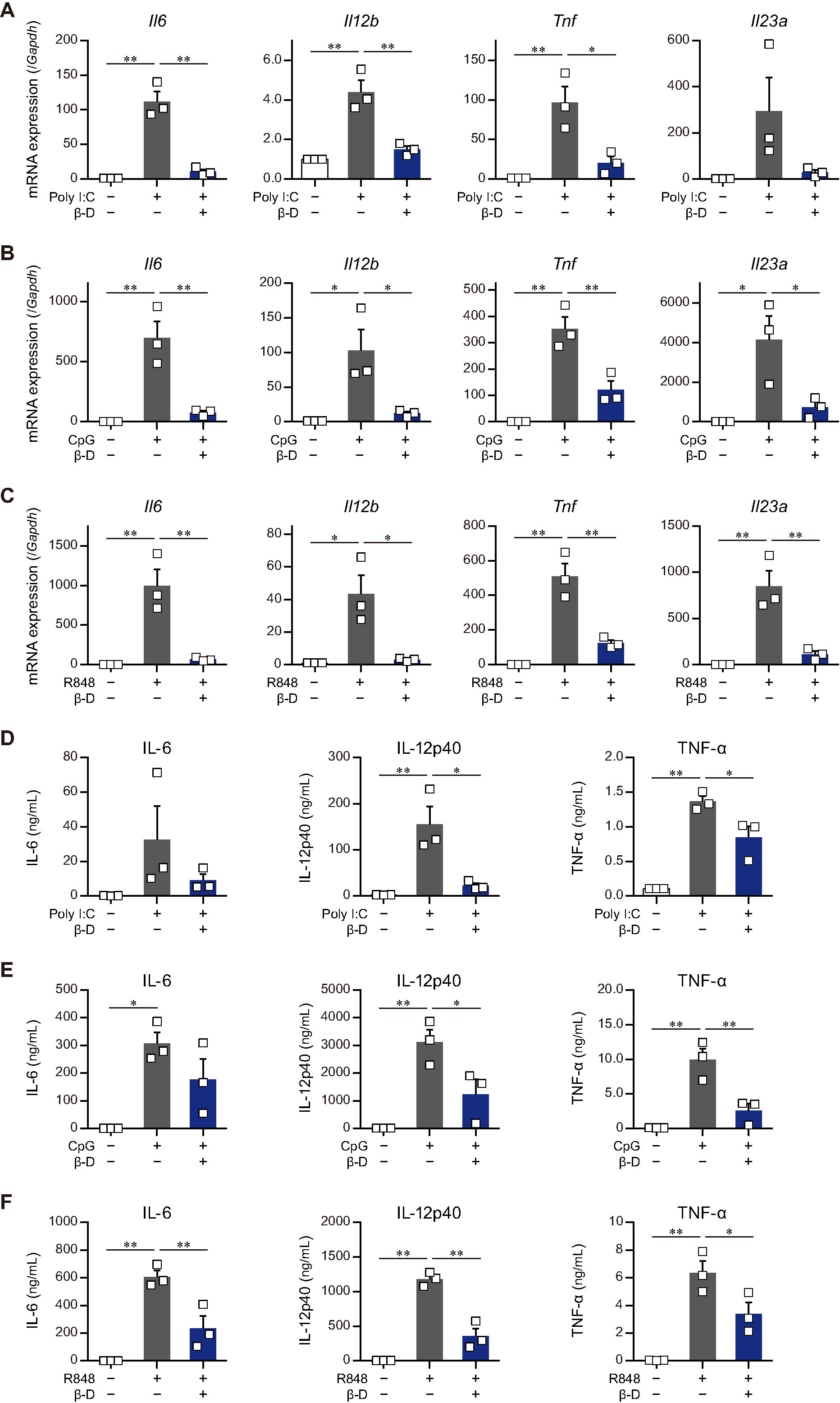
Suppressive effects of β-damascone on the stimulation of DCs by various TLR ligands. The effects of β-damascone (50 μM) on mRNA expression levels in (**A**, **B**, and **C**) and the protein release of cytokines (**D**, **E**, **F**) from BMDCs stimulated with polyI:C (**A** and **D**), CpG (**B** and **E**), or R848 (**C** and **F**). Data represent the mean ± SEM of three independent experiments performed in triplicate samples. The Tukey-Kramer test was used. *; *p* < 0.05, **; *p* < 0/01.

These results suggest that β-damascone exhibited suppressive effects on the TLR ligand-induced activation of DCs.

### Involvement of the NRF2 pathway in suppressive effects of β-damascone on DCs

Some substances derived from plants, particularly phytochemicals (e.g., sulforaphane), activate the NRF2 pathway by directly acting on Keap1, resulting in the induction of antioxidant and anti-inflammatory responses (Kensler et al. 2013). To clarify whether β-damascone activates the NRF2 pathway in DCs, we determined the NRF2 protein and *Hmox1* mRNA levels in β-damascone-treated DCs. As shown in **Figure 4A**, the Western blot analysis revealed that NRF2 protein levels increased in DCs in the presence of β-damascone and peaked 1 h after the addition of β-damascone to the culture medium. The mRNA levels of *Hmox1*, a target gene of NRF2, were markedly higher after the treatment with β-damascone than with LPS (**Figure 4B**). These increases in NRF2 protein and *Hmox1* mRNA levels in β-damascone-treated DCs suggest that β-damascone activated the NRF2 pathway in DCs. We then investigated the roles of NRF2 in the β-damascone-mediated modification of DC functions using *Nrf2^−/−^* DCs. We confirmed that β-damascone-induced increases in *Hmox1* mRNA levels in DCs were mostly abolished by a NRF2 deficiency (**Figure 4C**). When OVA-pulsed *Nrf2^−/−^* BMDCs were cocultured with OT-II CD4^+^ T cells under Th1-polarizing conditions, the suppression of Th1 development by β-damascone was not observed, whereas control (*Nrf2^+/−^*) DC-dependent Th1 development was significantly inhibited in the presence of β-damascone (**Figure 4D**), as was the case for WT BMDCs (**Figure 1D**). Furthermore, IL-12p40 release from LPS-stimulated DCs tended to increase by *Nrf2* deficiency in DCs, and the suppressive effect of β-damascone on IL-12p40 production was markedly reduced in *Nrf2^−/−^* BMDCs compared with that in *Nrf2^+/−^* BMDCs (**Figure 4E**).

**Figure 4.**
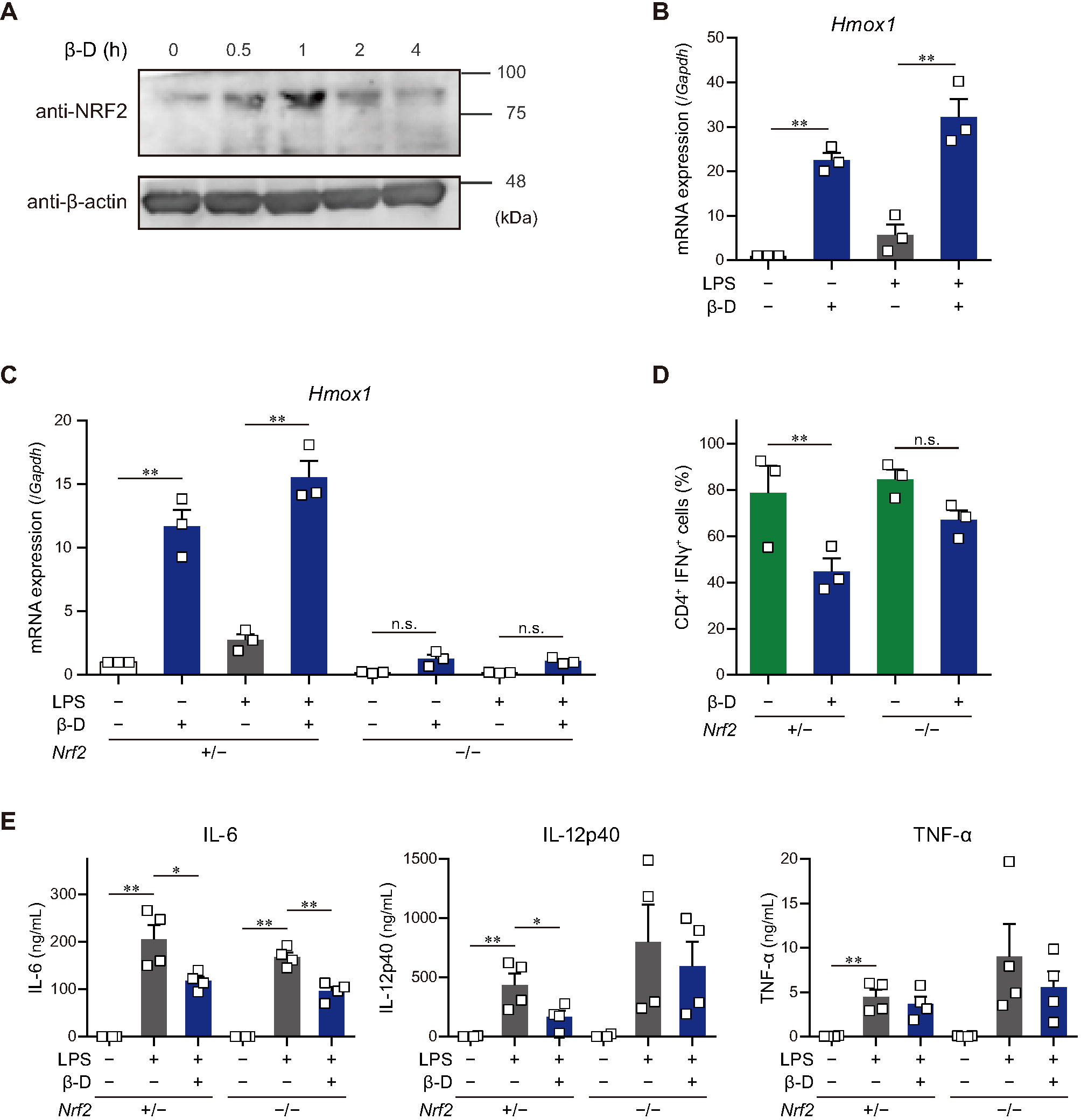
β-Damascone activated the NRF2 signaling in DCs, and a NRF2 deficiency attenuated its suppressive effects. **A**. A Western blot profile showing NRF2 protein levels in BMDCs following a treatment with β-damascone. BMDCs were incubated in the presence of 50 μM β-damascone for the indicated times. **B**. *Hmox1* mRNA levels in BMDCs. BMDCs cultured in the presence or absence of 50μM β-damascone for 24 h were incubated with or without 100 μg/mL LPS for an additional 3 h. **C**. *Hmox1* mRNA levels in BMDCs generated from NRF2-deficient mice (*Nrf2*^−/−^) and control mice (*Nrf2^+^*^/-^). **D**. Th1 development induced by a coculture with *Nrf2*^−/−^ BMDCs or control BMDCs. **E**. LPS-induced cytokine production by *Nrf2*^−/−^ BMDCs and control BMDCs. Data represent the mean ± SEM of more than three independent experiments performed in triplicate. The Tukey-Kramer test was used. *; *p* < 0.05, **; *p* < 0/01, n.s.; not significant.

These results demonstrate that DC functions, including Th1 induction and IL-12 production, were suppressed by β-damascone in a manner that was dependent on the activation of the NRF2 pathway in DCs.

### Oral administration of β-damascone ameliorated CHS in mice

We investigated whether β-damascone, which inhibited DC-mediated immune responses in *in vitro* experiments, exerted biologically significant effects on immune-mediated diseases using CHS model mice (**Figure 5A**). We examined the effects of various doses of β-damascone on ear swelling symptoms in DNFB-treated mice and found that the oral administration of β-damascone at 10 or 50 mg/kg per day from 2 days before sensitization suppressed ear swelling (data not shown). Under optimized doses, the inhibitory effects of β-damascone on ear swelling were observed not only after the 1st challenge, but also after the 2nd challenge (**Figure 5B**).

**Figure 5.**
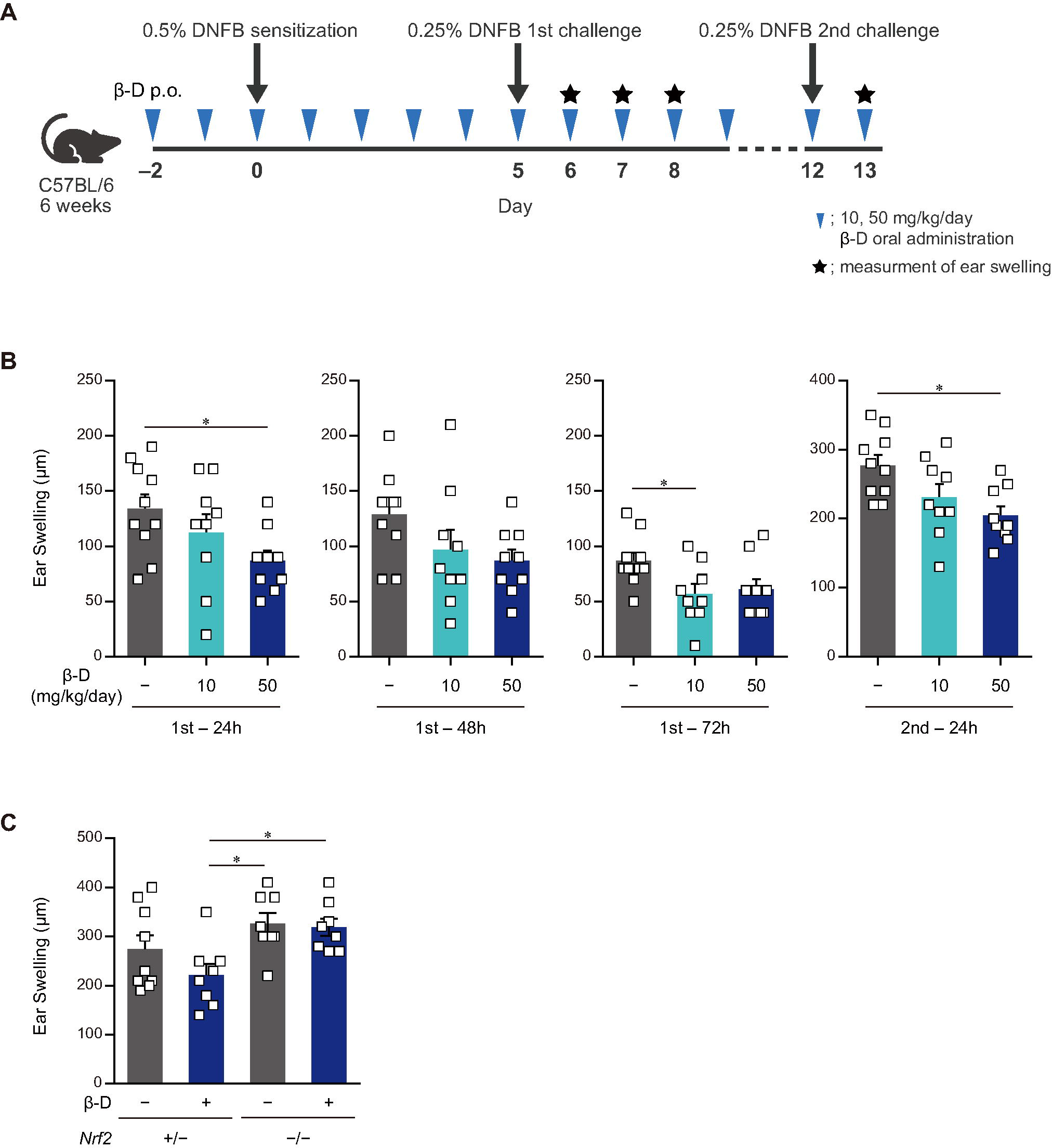
Orally administered β-damascone ameliorated the pathology of CHS. **A**. Schematic of the oral administration schedule of β-damascone in the DNFB-induced CHS model. The indicated amounts of β-damascone in 200 μl of saline were orally administered every day. p.o.; per os. **B**. Ear swelling in CHS-induced wild-type mice at the indicated time points. **C**. Ear swelling in CHS-induced *Nrf2* gene targeted mice 48 h after 2nd challenge. (Ear swelling) = (ear thickness after the challenge) – (ear thickness before the first challenge) The Tukey-Kramer test was used. *; *p* < 0.05.

We then utilized the CHS model in *Nrf2*^−/−^ mice to reveal the involvement of NRF2 in the protective effects of β-damascone *in vivo*. Ear swelling was tended to be more severe in *Nrf2*^−/−^ mice that in *Nrf2*^+/−^, and was not inhibited by the administration of β-damascone (**Figure 5C**).

These results indicate that orally administered β-damascone ameliorated CHS by modulating the function of NRF2 in mice.

## Discussion

In the present study, we identified β-damascone from among approximately 150 compounds as a novel immunomodulator through the 2-step screening and monitoring of DC-mediated immune responses. In *in vivo* experiments using a mouse model, we demonstrated that the oral intake of β-damascone, which activates the NRF2 pathway in DCs, attenuated CHS inflammation. NRF2 functions as a master regulator of antioxidant and anti-electrophilic responses by transactivating the genes encoding antioxidant enzymes and drug-metabolizing enzymes. In a previous study using a 2,4-dinitrochlorobenzene (DNCB)-induced CHS model, NRF2-deficient mice exhibited more severe inflammation accompanied by increased chemokine production and neutrophil recruitment in the skin than control mice (Helou et al. 2019). This finding showing that a NRF2 deficiency exacerbated CHS was consistent with the present results, which indicates important roles for NRF2 in DC-mediated immune responses. In addition to CHS, various immune-mediated inflammatory diseases and autoimmune diseases, including colitis and psoriasis, were shown to be exacerbated in NRF2 knockout mice (Khor et al. 2006, Ogawa et al. 2020, Yoh et al. 2001, Johnson et al. 2010). Although the effects of a NRF2 deficiency are not restricted to DC functions because NRF2 is ubiquitously expressed, the findings of these studies using NRF2-deficient mice support the potential of β-damascone for the prevention and/or treatment of inflammatory diseases and autoimmune diseases. We demonstrated that β-damascone significantly suppressed TLR7-mediated gene expression in and the protein production of IL-6, IL-12p40, and TNF-α by BMDCs (**Figure 3C** and **3F**). Since the topical application of imiquimod (Schön and Schön 2007, Gilliet et al. 2004), a ligand of TLR7 and TLR8, has been shown to causes a psoriasis-like pathology in mouse skin through the activation of dermal DCs and Langerhans cells (van der Fits et al. 2009), we expect β-damascone to ameliorate psoriasis, even from the viewpoint of modifications to DC functions. We intend to examine the effects of β-damascone on various immunorelated diseases, such as psoriasis, inflammatory bowel disease, and multiple sclerosis, using a mouse model in the near future.

Although β-damascone inhibited the DC-mediated development of Th1 cells from naïve CD4^+^ T cells (**Figure 1D**), the APC-independent development of Th1 cells by the stimulation of naïve CD4^+^ T cells with plate-coated anti-CD3 and anti-CD28 Abs in Th1-poralizing condition was not affected by β-damascone (data not shown). These results suggest that β-damascone suppressed Th1 development by modulating DC functions, which may involve the reductions in the release of IL-12 and the cell surface expression of MHC II and CD86 in β-damascone-treated DCs (**Figure 2**). In *in vivo* experiments, the administration of β-damascone was initiated 2 days before the sensitization to prevent CHS by inhibiting the activation of DCs. However, we cannot exclude a possibility that β-damascone have directly suppressed the functions of Th1 cells and CD8^+^ T cells, which contributed to the amelioration of CHS.

The results obtained using *Nrf2*^−/−^ DCs indicated that the inhibitory effects of β-damascone on IL-12 production by DCs and the Th1 development activity of DCs were dependent on NRF2. In contrast, the suppressive effects of β-damascone on the production of TNF-α and IL-6 were still observed in *Nrf2*^−/−^ DCs, suggesting that β-damascone modulated DC functions through target(s) other than NRF2. A previous study reported that the induction of the proinflammatory cytokine genes, *Il1a*, *Il1b*, and *Il6* was inhibited by NRF2, which bound to these genes and subsequently suppressed the recruitment of RNA polymerase II to the target genes in macrophages (Kobayashi et al. 2016). However, under our experimental conditions, NRF2 did not contribute to the inhibition of IL-6 and TNF-α production by DCs. A more detailed analysis to identify other target(s) of β-damascone will lead to the development of novel anti-inflammatory immunomodulators.

The present results showed that β-damascone was the most effective compound from an aroma library and confirmed that its oral administration ameliorated CHS. Although we obtained β-damascone as a natural compound candidate of immunomodulators in the present study, further improvements are possible; briefly, β-damascone may be developed as a drug that exerts stronger effects with less cytotoxicity through chemical modifications to its structure. Further research using an organic chemical approach with β-damascone as a lead compound is needed to develop a novel immunomodulators.

## Supporting information

Supplementary Table SI, SII, Figure S1, S2, and S3

## Abbreviations

Ab: antibody
Ag: antigen
APCs: antigen-presenting cells
BMDCs: bone marrow-derived dendritic cells
CHS: contact hypersensitivity
DCs: dendritic cells
DNFB: 2,3-dinitrofluorobenzene
NRF2: NF-E2-related factor 2
OVA: ovalbumin
p.o.: per os

## Authorship Contribution

Contribution: N.K. performed experiments, analyzed data, and wrote the paper; H.O. performed experiments and analyzed data; M.H. analyzed data and wrote the paper; M.A. N.I. K.N., Y.Y., I.H., and T.Y. performed experiments; G.I. and M.Y. provided experimental tools; C.N. designed research and wrote the paper.

## Acknowledgments

We thank the members of the Laboratory of Molecular Biology and Immunology, Department of Biological Science and Technology, Tokyo University of Science for constructive discussions and technical support.

