## Supplementary Table SI, SII, Figure S1, S2, and S3 for "A rose flavor compound activating the NRF2 pathway in dendritic cells ameliorates contact hypersensitivity in mice": Kodama-Okada-Front Nutr-Supplementary Information.docx

**Supplemental Information**

**Supplementary Table SI.** The information of primers used in quantitative PCR.

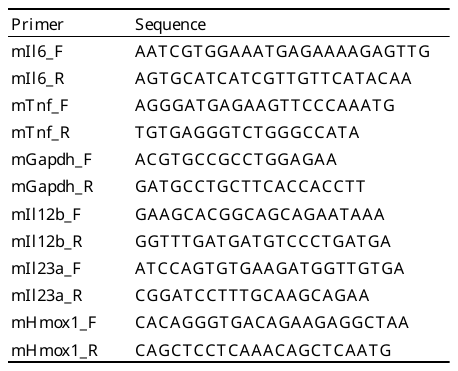

**Supplementary Table SII.** A list of compounds in an aroma chemical library.

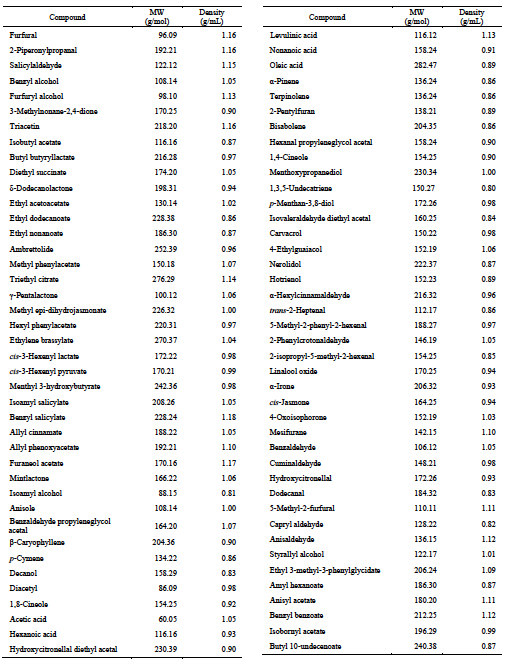

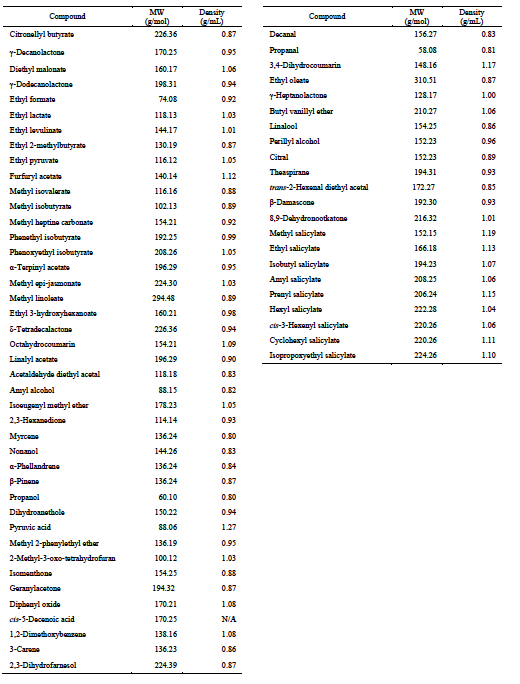

**Supplementary Figures and Figure Legends**

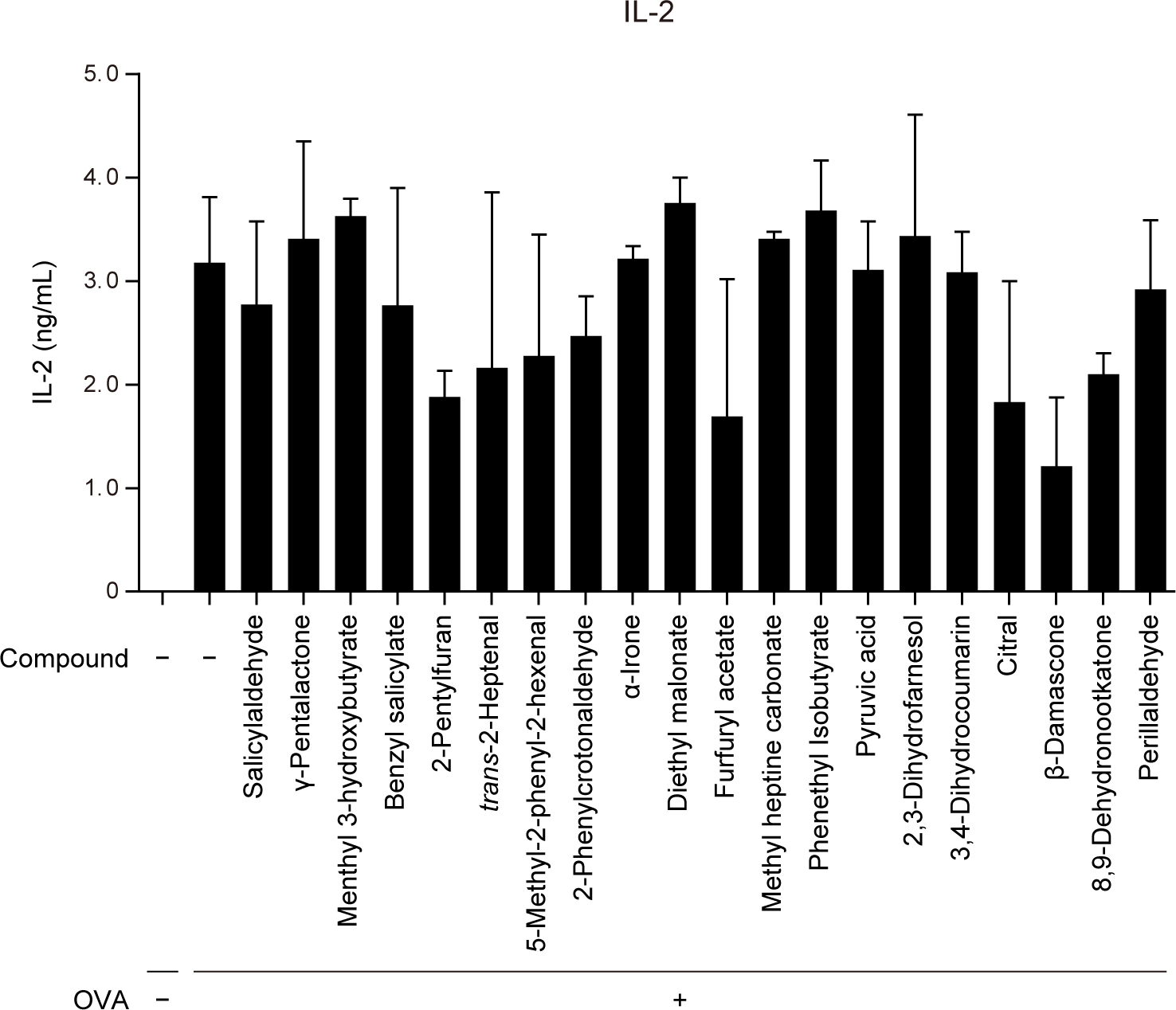

**Figure S1.** IL-2 production levels in the second screening.

The numbers of compounds are those in a library list (**Supplementary Table SII**). Each compound was added to the culture media of OVA-pulsed OT-II spleen cells at 0.001% (vol/vol) of the final concentration.

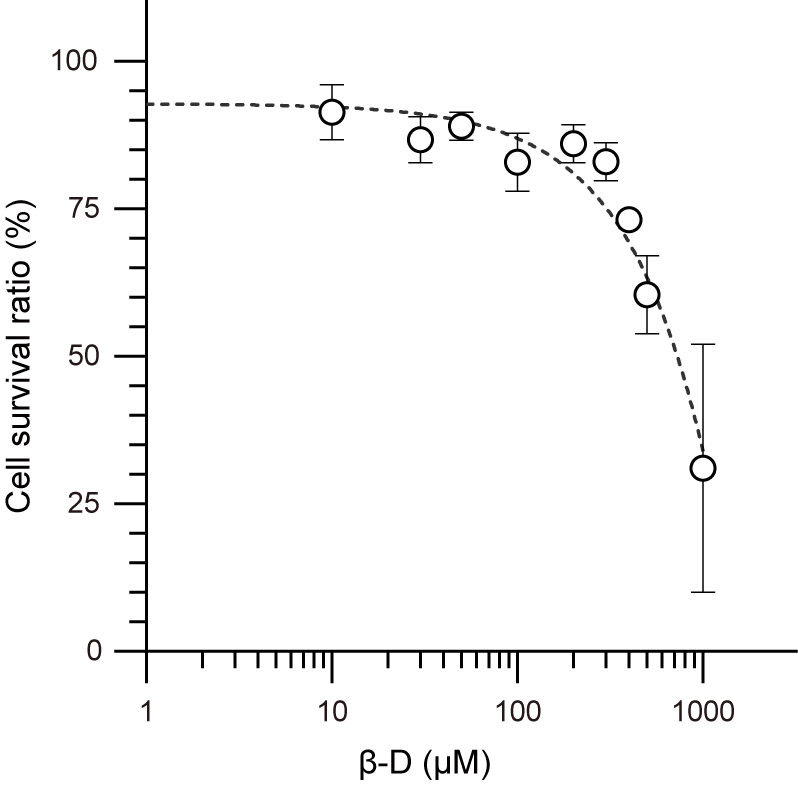

**Figure S2.** Cell viabilities of DCs in the presence of β-damascone.

BMDCs were preincubated with the indicated concentrations of β-damascone for 24 h. Cell viability was judged with DAPI staining.

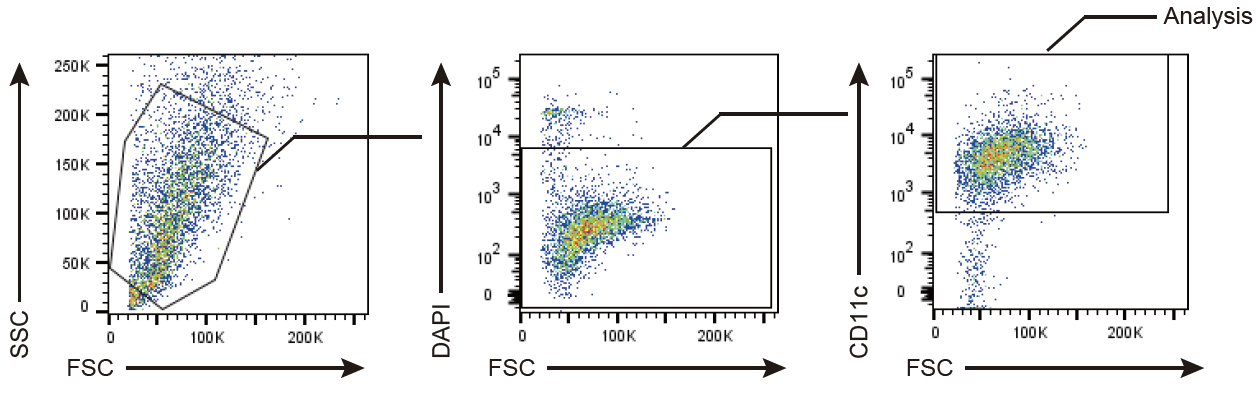

**Figure S3.** Gating strategies of flow cytometric analyses of BMDCs.

DAPI^-^/CD11c^+^ population in BMDCs was gated to determine the expression levels of MHC class II and CD86.
